# Across Species Identification of Genes Bridging Cognition and Reproduction

**DOI:** 10.64898/2026.07.02.736122

**Authors:** Zeynep Kizilaslan, Jessica Townsend Graybeal, Christopher Huffman, Andres Mejia Montoya, Francisco Peñagaricano, Mehmet Kizilaslan, Nagib Ahsan, Hasan Khatib

## Abstract

Evolutionary success in mammals requires coordinated regulation of cognitive functions and reproductive capacity. Such coordination must involve shared genes and molecular pathways between the brain and germ cells, yet direct evidence linking cognition to reproduction across species remains limited. Here, proteomic and transcriptomic analyses were performed experimentally in *Ovis aries* and *Rattus norvegicus*, while transcriptomic datasets from *Mus musculus*, *Macaca mulatta*, and *Homo sapiens* were analyzed in silico. We identified 8,464 protein-coding genes shared between the brain and sperm/testis and conserved across five species. In rats, 8,444 of these genes were also shared between the brain and the ovary. Functional annotation classified 3,890 genes as associated with both neurological and reproductive functions, and 1,752 as uncharacterized in these contexts, highlighting candidates for future studies on reproductive and neurological disorders. These findings reveal a deeply conserved genetic network linking neurological and reproductive systems, underscoring the evolutionary interplay that supports mammalian fitness.

## INTRODUCTION

The evolution of complexity in organisms depends on coordinated adaptations among organ systems that shape fitness-related traits, allowing multiple developmental mechanisms and networks to coevolve without reducing overall fitness ^1^. Among these mechanisms, the neurological and cognitive processes that enable organisms to perceive, memorize, interpret, and respond to environmental stimuli have been linked to fitness advantages. Indeed, the correlation between cognition and reproductive success has been investigated in birds ^2–4^. Cole et al.^2^ found that problem-solving ability was positively associated with clutch size and the number of fledglings in wild great tits. In contrast, evidence connecting cognition to reproductive fitness in mammals remains scarce. To date, only one study has reported positive correlations between intelligence and semen quality traits—sperm concentration, count, and motility—in 425 U.S. Army veterans ^5^.

Pleiotropy plays a central role in coordinating biological systems such as cognition and reproduction by allowing the same genes to act across molecular pathways that affect multiple phenotypic traits ^6,7^. Examples of pleiotropic effects include mutations in the *HTT* gene, which cause Huntington’s disease, an inherited neurodegenerative disorder, and lead to smaller testes, azoospermia, and male infertility in mice ^8^. Another example is the *FMR1* gene, responsible for fragile X syndrome, the most common inherited cause of intellectual disability. Mutations in *FMR1* are also associated with ovarian dysfunction and diminished ovarian reserve in women and have been shown to affect Sertoli cell proliferation during testicular development in mice ^9–11^. Thus, single-gene pleiotropic relationships between neurological and reproductive functions have been documented, yet the overall extent of the connection between the brain and reproductive systems remains poorly understood.

Recent studies suggest substantial overlaps in genes and proteins between the brain and the testis ^12,13^. For example, a comparative transcriptomic study across 17 human tissues using 760 Unigene datasets found the highest similarity between the brain and testis, with 364 genes (48%) commonly expressed in both tissues ^12^, suggesting shared regulatory pathways. Proteomic analyses further support this overlap, identifying 13,442 proteins common to both the brain and testis, accounting for 14,315 and 15,687 proteins, respectively, many of which are involved in exocytosis, tissue formation, and neuronal processes ^13^. However, these transcriptomic and proteomic studies were analyzed in silico. Furthermore, functional studies corroborate these molecular parallels: mammalian sperm and neurons express similar neuronal receptors, including GABAA receptors, which regulate neuronal excitability and are also present in sheep, pig, and human sperm membranes ^14–16^. GABAA receptors have been shown to modulate key sperm functions, including motility, capacitation, hyperactivation, and the acrosome reaction ^17^. Interestingly, recently we found that the majority of sperm genes in sheep affected epigenetically by paternal diet have functions in the nervous system and brain development, neuron projection, and synaptic signaling ^18–21^.

Collectively, these transcriptomic, proteomic, and functional studies reveal extensive molecular and functional overlaps between the brain and reproductive tissues, highlighting the need to better understand the mechanisms linking neurological and reproductive traits. However, no study to date has examined the evolutionary conservation of the brain–testis transcriptional relationship across multiple species, nor has it investigated how this relationship compares with that between the brain and the ovaries. To address these gaps, we hypothesize that evolutionary success depends on coordination between cognitive adaptation, driven by the brain, and reproductive fitness, mediated by sperm and ovaries. Accordingly, this study investigated the extent of shared gene and protein expression among the brain, testis, sperm, and ovary, integrating experimental data from *Ovis aries* and *Rattus norvegicus* with in silico analyses in *Mus musculus*, *Macaca mulatta*, and *Homo sapiens*.

## RESULTS

### Brain and sperm show high similarity in protein and gene expression in sheep and rats

To identify shared proteins and transcripts, we conducted integrated proteomic and transcriptomic analyses of 10 distinct brain regions, sperm, and white blood cells (WBCs) collected from five male sheep (Fig. 1). Using a bottom-up proteomics approach, we quantified 4,072 annotated proteins across these tissues. The 10 brain regions exhibited highly similar protein expression patterns (Fig. 1A, B), prompting us to combine them into a composite brain dataset for comparison with sperm and WBC profiles. Among the detected proteins (Supplementary File 1, Table S1), 2,493 (61.2%) were shared between brain and sperm, while 2,717 (66.7%) were shared between brain and WBC.

**Figure 1.**
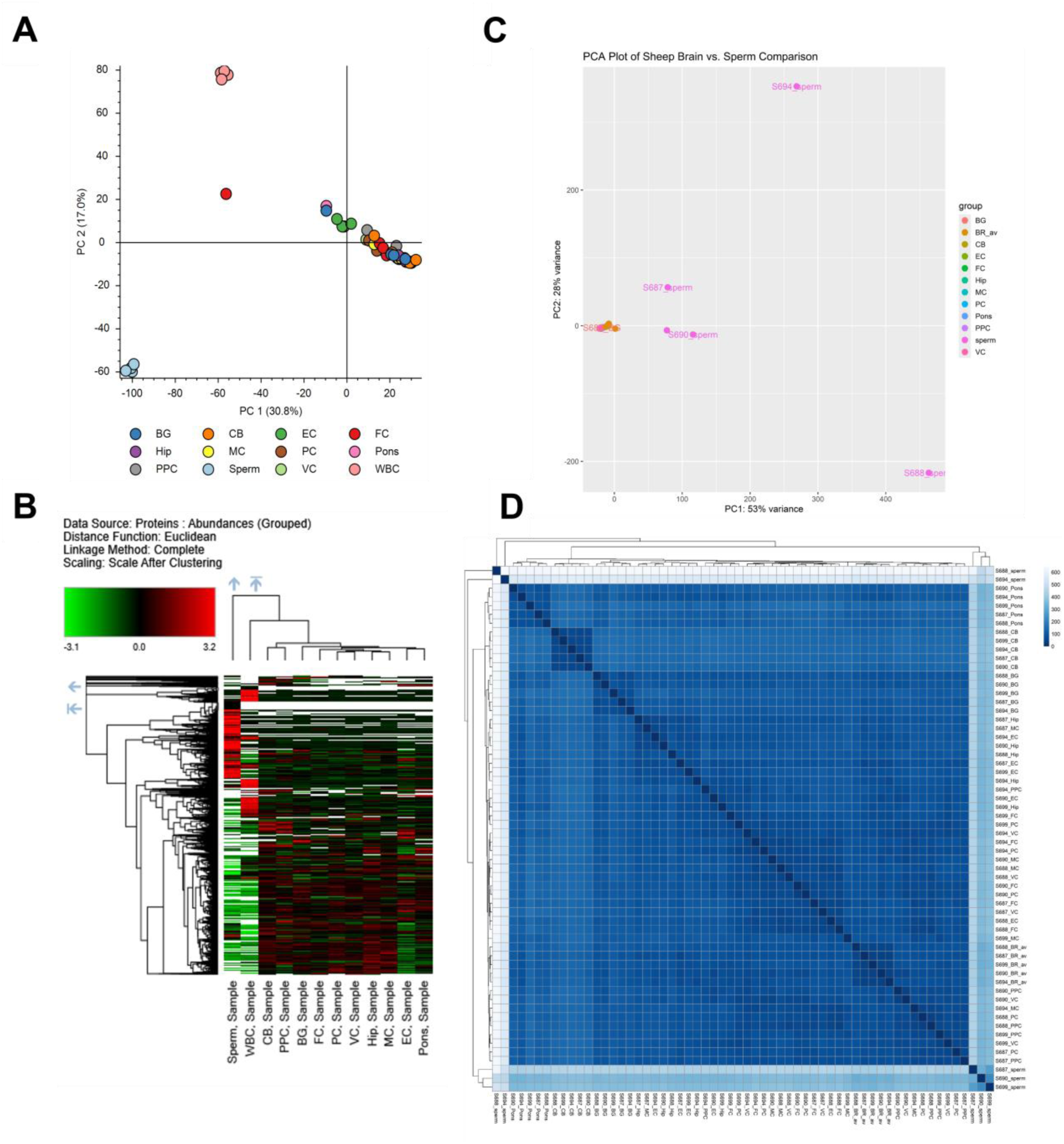
Proteomic and transcriptomic profiles of brain regions, sperm, and WBC in sheep. (A) PCA of proteomic profiles from ten brain regions, sperm, and WBC. (B) Heatmap of proteomic abundances in ten brain regions and sperm. (C) PCA of RNA-Seq expression profiles from brain dissections and sperm based on normalized read counts. (D) Heatmap of log-transformed, normalized transcript counts showing sample-level expression similarities.

We next examined gene expression similarity between each of the 10 brain regions and sperm. Of the 31,304 annotated sheep genes, we identified region-specific expression in 3,651 genes in basal ganglia (BG), 2,431 in cerebellum (CB), 4,026 in entorhinal cortex (EC), 3,741 in frontal cortex (FC), 4,045 in hippocampus (Hip), 3,713 in motor cortex (MC), 3,813 in prefrontal cortex (PC), 3,887 in pons, 3,764 in posterior parietal cortex (PPC), and 3,800 in visual cortex (VC) compared to sperm. Given the strong similarity across brain regions, we computed a brain average (BR_av) expression profile to compare with sperm (Fig. 1C, D).

Out of 21,794 expressed genes, 14,955 (68.6%) showed shared expression between at least one brain region and sperm and were used for downstream analyses (Supplementary File 1, Table S2). In contrast, 4,933 genes were brain-specific, 1,906 were sperm-specific, and 9,510 showed no expression in either tissue. All brain regions, including the BR_av composite, clustered tightly together, reflecting their high expression similarity, whereas sperm samples showed somewhat greater variability in their profiles (Fig. 1C, D). The first two principal components explained 81% of the total variance in gene expression across tissues (Fig. 1C), highlighting high concordance among brain regions and a moderately high, yet distinct, similarity between sperm and brain samples, as further supported by the sample heatmap (Fig. 1D).

We extended our experimental investigation to rats to determine whether the observed pattern of shared gene expression between whole brain and sperm is specific to a single species or reproducible across two species from different taxonomic suborders. Among the 22,281 expressed genes identified in rats, 16,072 (72.1%) were shared between brain and sperm, 4,503 (20.2%) were brain-specific, and 1,706 (7.3%) were sperm-specific (Supplementary File 1, Table S3).

### Shared genes between brain, sperm, and testis are evolutionarily conserved

To test our hypothesis that genes shared between the brain and sperm/testis are evolutionarily conserved across mammals, we expanded our analysis from *Ovis aries* (sheep) and *Rattus norvegicus* (rat) to include *Mus musculus* (mouse), *Macaca mulatta* (Rhesus macaque), and *Homo sapiens* (human). For mouse, rhesus macaque, and human, we conducted *in silico* analyses using transcriptomic data from brain and sperm/testis tissues obtained from three public databases. Remarkably, a total of 8,464 protein-coding genes showed shared expression between brain, sperm, and testis across all five species (Fig. 2A; Supplementary File 2, Table S4). Among them, humans exhibited the highest number of species-specific brain–testis shared genes (n = 6,604), while mice showed the lowest number of species-specific brain–testis shared genes (n = 1,406) (Fig. 2A). Figure 2B summarizes the total number of genes analyzed, tissue-specific genes, and the number of shared genes between brain and sperm/testis for each species. The largest number of shared brain–testis genes was observed in humans (n = 20,935), whereas the smallest was found in mice (n = 13,680) (Supplementary File 2, Table S4).

**Fig. 2.**
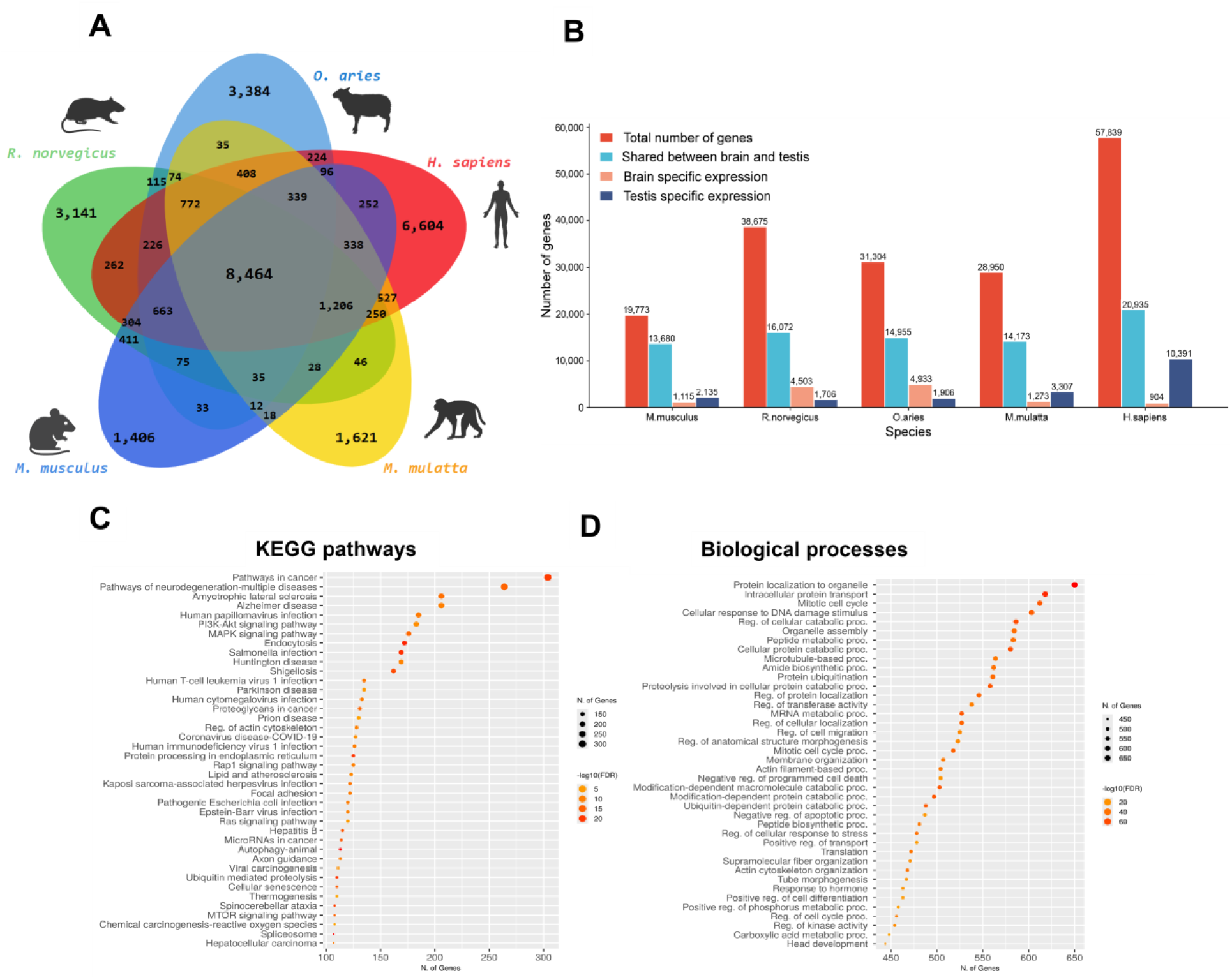
Expression profiles, KEGG pathways, and biological functions of genes shared between brain, sperm, and testis across five species. (A) Venn diagram showing genes shared between brain, sperm, and testis in *Ovis aries*, *Homo sapiens*, *Macaca mulatta*, *Mus musculus*, and *Rattus norvegicus*. (B) Pairwise comparisons of shared genes between species. In *O. aries* and *R. norvegicus*, shared genes are between the brain and sperm, whereas in the other three species, shared genes are between the brain and the testis. (C) KEGG pathway enrichment of the 8,464 shared genes across species. (D) Biological process annotation of brain–testis shared genes across all five species

There were also significant pairwise overlaps in gene expression between species, with the highest observed between Rhesus macaque and human (18,810 overlapping genes between the brain and testis) and the lowest between sheep and mouse (9,717 overlapping genes between sheep brain–sperm and mouse brain–testis (Fig. 2A; Supplementary File 2, Table S5). When considering both shared and tissue-specific genes (brain-, sperm-, or testis-specific), the difference between sperm and testis was minimal, indicating a high degree of similarity in their gene expression profiles.

To understand the biological significance of cross-species-conserved genes shared between the brain and sperm/testis, we performed functional enrichment analysis to identify key KEGG pathways and biological processes associated with these genes (Fig. 2C, D). The results revealed a strong enrichment of genes involved in neurological functions, with three of the top four KEGG pathways (neurodegeneration–multiple diseases, amyotrophic lateral sclerosis, and Alzheimer’s disease) related to neurodegeneration. Each of these pathways included more than 200 genes from our dataset of conserved shared genes across species (Fig. 2C). In addition, many enriched KEGG pathways and biological processes were associated with neurodevelopment, immune response, and reproduction, as well as diseases related to these systems (Fig. 2C, D). For instance, 276 of the 8,464 genes shared between brain and testis across species were mapped to the neurodegeneration pathway (hsa05022) in KEGG, which contains 367 annotated genes (see Supplementary Fig. S1).

### Expression levels of shared brain and sperm/testis genes are conserved across species

To determine whether the evolutionarily conserved genes shared between brain and sperm/testis also exhibit conserved expression patterns, we performed differential expression analyses for each species, comparing brain and sperm/testis transcriptomes of the 8,464 shared genes. Using thresholds of ≥10-fold change and FDR-corrected *q* < 0.05, we identified 7,954 genes in sheep, 8,012 in rat, 7,953 in mouse, 8,138 in rhesus macaque, and 8,007 in human that showed similar expression levels between brain and sperm/testis samples (Fig. 3A). Overall, 7,061 genes demonstrated comparable expression across all five species (Supplementary File 2, Table S4).

**Fig. 3.**
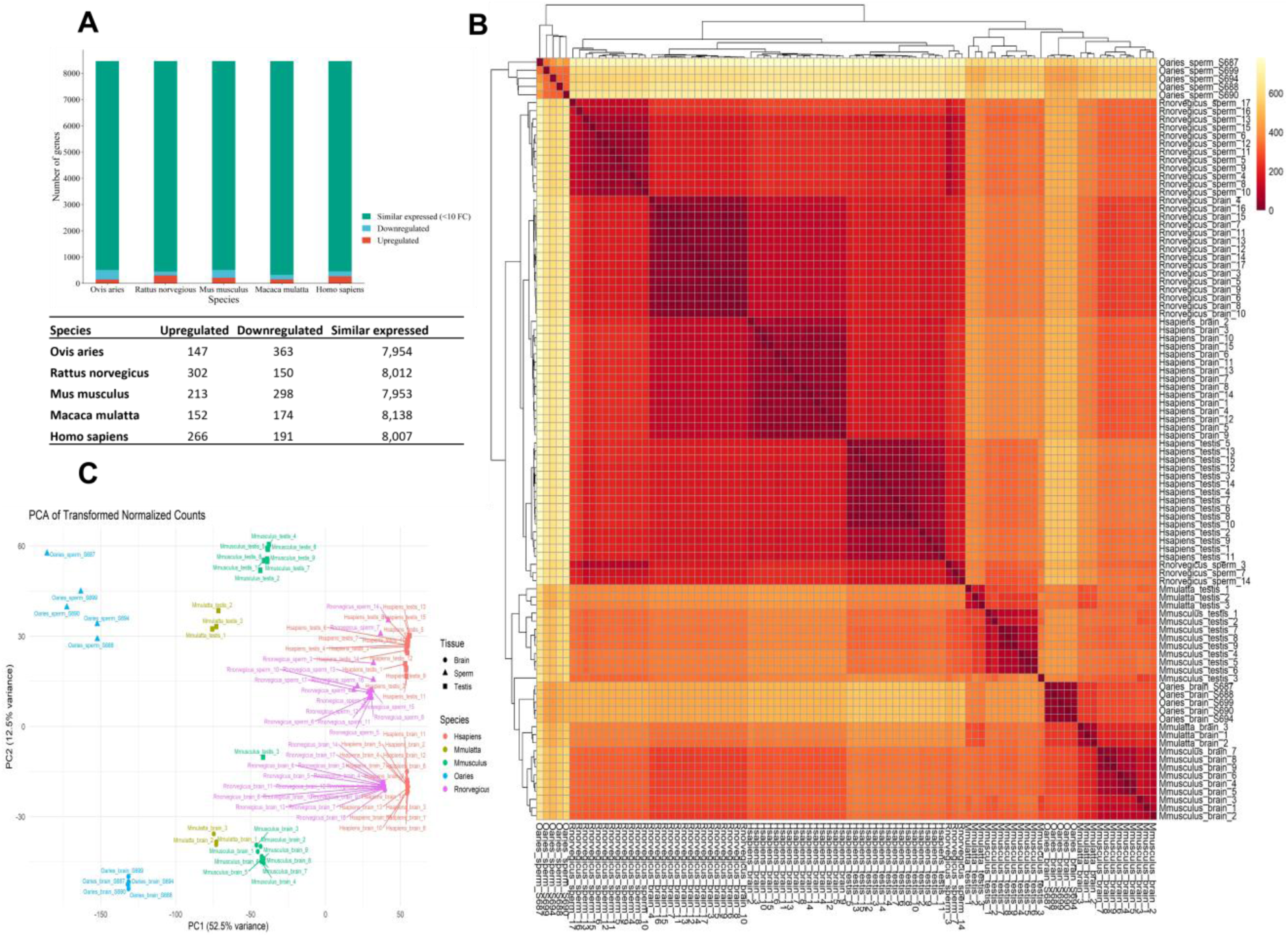
Differential gene expression analysis of brain–sperm/testis shared genes across five species. (A) Number of genes with similar expression (<10-fold change), upregulated, or downregulated in each species. (B) Across-species heatmap of normalized expression profiles showing sample-wise similarities among brain–sperm/testis shared genes. (C) PCA of expression profiles for 8,464 genes shared between brain and sperm/testis across all five species.

Only a small number of genes were differentially expressed between the brain and sperm/testis within each species. Specifically, there were 147 upregulated and 363 downregulated genes in sheep sperm, 302 up and 150 down in rat sperm, 213 up and 298 down in mouse testis, 152 up and 174 down in Rhesus macaque testis, and 266 up and 191 down in human testis (Fig. 3A). The overlaps of differentially expressed genes (DEGs) among species are summarized in Table 2, revealing only nine genes that showed consistent cross-species differential expression between brain and sperm/testis. Consistent with these results, the hierarchical clustering heatmap (Fig. 3B) shows high similarity in the expression profiles of shared genes between brain and sperm/testis across species. The strongest similarity was observed between rat brain–sperm and human brain–testis samples, differing by only ∼200 genes. This was followed by Rhesus macaque and mouse brain–testis samples, whereas sheep sperm samples showed the greatest divergence, differing by approximately 600 genes.

A similar pattern emerged from the principal component analysis (PCA; Fig. 3C). The first principal component (PC1), which explained 52.5% of the variance, grouped human, rat, mouse, and Rhesus macaque samples closely together, whereas sheep samples were slightly separated from the cluster. The second principal component (PC2) accounted for only 12.5% of the variance, indicating relatively minor differences among the species along this axis.

### Comparative analysis reveals high conservation between brain–ovary and brain–sperm genes across species

To further test our hypothesis that evolutionary success relies on coordinated function between the nervous and reproductive systems, we analyzed the extent of shared gene expression between brain and ovary tissues in 14 female rats using RNA-Seq. We identified 17,794 shared genes between brain and ovary, along with 1,931 ovary-specific and 2,705 brain-specific genes (Fig. 4A; Supplementary File 3, Table S6). We next examined whether these rat brain–ovary shared genes overlap with the brain–sperm shared genes conserved across the five species. Remarkably, 8,444 of the 8,464 evolutionarily conserved brain–sperm shared genes were also present among the rat brain–ovary shared genes (Fig. 4B; Supplementary File 3, Table S6). Further analysis revealed substantial overlaps between rat brain–ovary shared genes and the brain–sperm/testis shared genes of other species: 10,799 with sheep, 15,518 with rat, 11,820 with mouse, 11,496 with Rhesus macaque, and 13,052 with human (Fig. 4B; Supplementary File 3, Table S6).

**Fig. 4.**
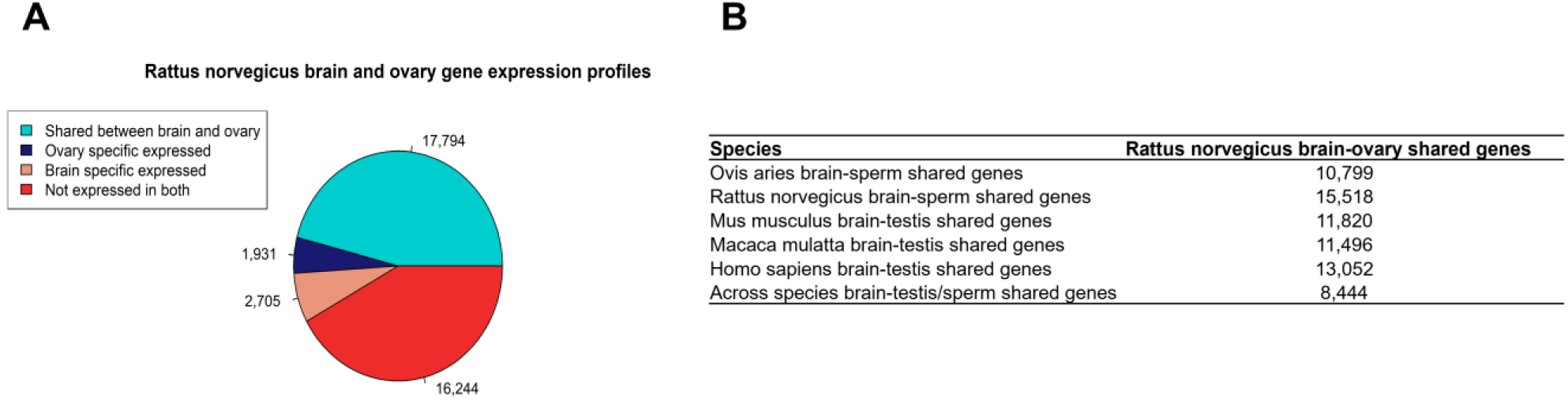
Comparison of shared genes between brain, sperm, testis, and ovary across five species. (A) Gene expression profiles of brain and ovary in *Rattus norvegicus*. (B) Overlap of 17,794 brain–ovary shared genes in rats with brain–sperm/testis shared genes conserved across all species.

### Integrative functional annotation of brain–testis genes across species highlights neurological and reproductive functions

To gain deeper insight into the functional roles of genes shared between the brain and the testis, we conducted a comprehensive functional annotation of the 8,464 cross-species conserved genes identified in this study. Using data integrated from 10 curated databases (Supplementary File 4), we classified these genes according to their associations with biological pathways, diseases, and processes (Fig. 5; Supplementary File 4, Table S7). Among the 8,464 genes, 2,289 were annotated with neurological functions, and 533 were associated with reproductive functions. Notably, 3,890 genes were linked to both neurological and reproductive pathways, diseases, or processes, underscoring their potential dual roles in these systems. The remaining 1,752 genes had no known functional annotations related to either the nervous or reproductive systems (Fig. 5; Supplementary File 4, Table S7).

**Fig. 5.**
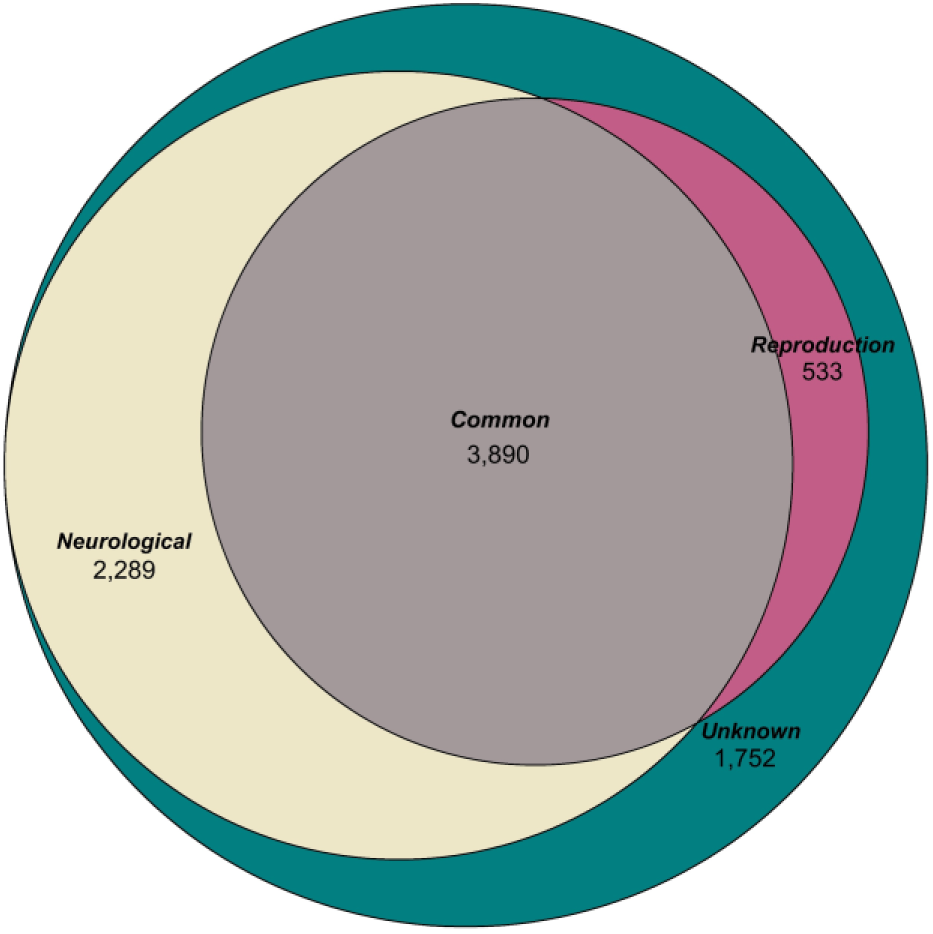
Functional annotation of 8,464 genes shared between the brain and testis across species. Annotations were compiled from ten databases and classified according to known neurological and reproductive functions.

## DISCUSSION

Achieving evolutionary success in mammals requires coordinated regulation of cognitive functions by the brain and reproductive capacity by the sperm and ovary. Previous *in silico* studies have suggested that the brain and testis share the greatest number of genes and proteins compared with other tissues in mice and humans ^12,13^. However, given the limitations of in silico analyses, we sought to experimentally determine the extent of shared genes and proteins in sheep and rats. Proteomic and transcriptomic analyses in sheep revealed high similarity in protein and mRNA expression among the 10 brain regions. Therefore, data from all brain regions were combined for comparative analysis with sperm. On average, the sheep brain shared 66.7% of its proteins and 68.6% of its transcripts with sperm. Rats showed slightly higher brain-sperm transcriptomic similarity (72.1%).

These findings prompted us to investigate the evolutionary conservation of shared gene expression across additional mammalian species. We retrieved RNA-Seq data for Rhesus macaque, mouse, and human from three independent transcriptomic databases, focusing on brain and testis tissues. For each species, we identified genes expressed in both tissues and intersected the resulting gene lists to identify those shared across species. In total, our analysis identified 8,464 protein-coding genes shared between brain and sperm/testis tissues across five species (Fig. 2A, B). Notably, the pathways containing the largest numbers of these shared genes were associated with cancer, several neurodegenerative disorders, amyotrophic lateral sclerosis (ALS), and Alzheimer’s disease (Fig. 2C).

A high proportion of these shared genes in each species exhibited remarkably similar expression levels between brain and sperm/testis samples. Specifically, 93.9% of the shared genes in sheep, 94.6% in rats, 93.9% in mice, 96% in macaques, and 95% in humans showed consistent expression similarity between brain and sperm/testis tissues. Overall, 83% of the shared genes displayed similar expression patterns across all five species (Fig. 3A). Sheep brain and sperm samples showed the fewest shared genes overlapping with the other four species, both in terms of gene number and expression similarity (Fig. 3B, C). This pattern may reflect the earlier evolutionary divergence of sheep within the mammalian lineage. Phylogenetic consensus places sheep within the Laurasiatheria superorder, which diverged from the Euarchontoglires lineage (rats, mice, macaques, and humans) approximately 90 million years ago ^22^.

A total of 17,794 genes were identified as being shared between brain and ovary tissues in rats (Fig. 4A). Remarkably, of the 8,464 genes shared between brain and sperm/testis and conserved across species, 8,444 were also detected in both brain and ovary tissues. These findings suggest that the observed relationship extends to both sexes, indicating a broader alignment between the nervous and reproductive systems, at least as evidenced in rats (Fig. 4B). A possible physiological explanation for this overlap may lie in the process of sexual differentiation of the reproductive tract. During early embryonic development, both male and female embryos possess undifferentiated gonads that can develop into either testes or ovaries. In subsequent developmental stages, hormonal cues direct these structures toward male or female reproductive fates ^23^.

Our comprehensive annotation analysis using ten databases, followed by functional classification, assigned the 8,464 genes shared between brain and sperm/testis to the following categories: ‘annotated with neurological functions only’ (27%), ‘annotated with reproductive functions only’ (6%), ‘annotated with neurological and reproductive functions’ (46%), and ‘uncharacterized for either function’ (21%) (Fig. 5). The genes annotated for both neurological and reproductive functions across the five species (n = 3,890) included well-known examples such as *HTT*, *FMR1*, and *CAMK2A*. Mutations in *HTT*, which cause Huntington’s disease, have also been associated with male infertility and arrested spermiogenesis in mice^8^. *FMR1* plays extensively characterized roles in Fragile X syndrome, neurodegenerative disorders such as ataxia and tremor, and autism spectrum disorder ^24,25^. *FMR1* mutations are linked to primary ovarian failure and have been proposed as genetic factors influencing reproductive decision-making ^24,26^. *CAMK2A* mediates calcium signaling and is essential for hippocampal long-term potentiation and spatial learning ^27^. Mutations in *CAMK2A* disrupt dendritic morphology and synaptic transmission, resulting in ASD-related behaviors and intellectual disability ^28^. In the reproductive system, *CAMK2A* has been implicated in preventing spontaneous acrosomal exocytosis in sperm and in mediating Ca²-dependent signaling during oocyte activation in mice ^29,30^.

We also identified 1,752 genes lacking known functional associations with neurological or reproductive processes, including Cluster of Differentiation 72 (*CD72*) and *TMEM9* Domain Family Member B (*TMEM9B*). The *CD72* gene is localized to the plasma membrane, where it participates in transmembrane signaling and negatively regulates B cell receptor signaling. Previous studies have shown that *CD72* plays a major role in systemic lupus erythematosus by promoting B-lymphocyte activation and modulating the tumor microenvironment ^31,32^. The Tmem9b protein is localized to the early endosome and lysosomal membranes and positively regulates canonical NF-κB signaling. Its expression has been associated with modulation of the tumor necrosis factor (TNF) signaling pathway and is essential for the production of proinflammatory cytokines in response to TNF, interleukin-1β, and toll-like receptor (TLR) ligands ^33^. Given that these 1,752 genes show shared expression in the brain and sperm/testis across the five species, we propose them as novel candidate genes potentially involved in neurological and reproductive functions. These findings support the notion that the 8,464 genes represent key molecular mechanisms underlying shared phenotypes between the nervous and reproductive systems across sheep, rats, mice, macaques, and humans.

Interestingly, many genes shared between brain and sperm/testis across species appear to be involved in immune functions, particularly in immune responses and autoimmune conditions (Fig. 2C). These genes are also shared along the brain–ovary axis in rats. Similar to nervous and reproductive systems, the mammalian immune system has undergone intense evolutionary refinement driven by natural selection, conferring survival and reproductive advantages through complex intersystem interactions. For instance, neural reflexes modulate immune responses to pathogens, and neural mechanisms influence the development and regulation of immunity ^34^. Moreover, a recent study comparing neural, immune, and reproductive cells revealed striking morphological, molecular, and structural similarities, suggesting a shared evolutionary origin and functional convergence during the transition from unicellular to multicellular life ^35^. Collectively, these observations point to an evolutionary interplay among the neurological, reproductive, and immune systems that may have co-evolved as an integrated regulatory network. We propose that the identified genes represent promising candidates for future studies investigating these interconnected systems.

This study provides the first integrated experimental and in silico evidence for the evolutionary conservation of genes shared between the brain and sperm/testis across five mammalian species, demonstrated at both the gene identity and expression levels. Comprehensive functional annotation revealed that many of these shared genes have pleiotropic roles in the nervous and reproductive systems, while others represent newly identified candidates with potential dual functions. Notably, most of the conserved genes also exhibited shared expression patterns between the brain and sperm/testis within each species and extended conservation along the brain–ovary axis in rats. These findings highlight a set of previously uncharacterized genes as promising candidates for further investigation into the molecular mechanisms underlying neural and reproductive functions, as well as potential targets for studies of disease etiology, diagnosis, and prognosis in both humans and animals. Importantly, the observation that many evolutionarily conserved brain–testis shared genes are involved in immune processes suggests an intricate evolutionary interplay among the neurological, reproductive, and immune systems.

### Limitations of the study

The current study includes experimental data from two species (sheep and rats) and in silico analyses from three additional species (mouse, rhesus macaque, and human). Future studies would benefit from increased sample sizes for each species, particularly for female brain and ovarian tissues. Moreover, to further elucidate expression patterns in neural and reproductive tissues and to strengthen inferences about evolutionary conservation across mammals, additional experimental data from a broader range of species are warranted.

## RESOURCE AVAILABILITY

### Lead contact

Requests for further information and resources should be directed to and will be fulfilled by the lead contact, Hasan Khatib

### Materials availability

This study did not generate new unique reagents.

### Data and code availability

The data of the study have been deposited in the National Center for Biotechnology Information (NCBI)’s Gene Expression Omnibus (GEO) and is accessible through accession number GSE316889.

All other data used in this study are already from public databases of NCBI GEO and Human Genotype-Tissue Expression (GTEx) Atlas.

## ACKNOWLEDGMENTS

This research was supported by the USDA/NIFA Grants No. 2023-67015-39527 and 2024-67015-42244 to H.K. N.A. gratefully acknowledging the initial funding support from the OU VPRP Office for establishing the Proteomics Core Facility. N. A. also acknowledges the contributions of the volunteer undergraduate and graduate students who participated in the proteomics work.

## AUTHOR CONTRIBUTIONS

Conceptualization, H.K. and F.P.; methodology, H.K.; investigation, Z.K., J.T.G., C.H., A.M.M., and N.A.; data curation, M.K.; formal analysis, N.A. and M.K; writing—original draft, Z.K., M.K., and H.K.; writing—review & editing, Z.K., J.T.G., C.H., A.M.M., F.P., M.K., N.A., and H.K.; funding acquisition, H.K.; supervision, H.K.

## DECLARATION OF GENERATIVE AI AND AI-ASSISTED TECHNOLOGIES

During the preparation of this work, we used generative AI ‘Microsoft Copilot’ for the classification of functional annotations obtained for the genes, which is also cited in the relevant section.

## SUPPLEMENTAL INFORMATION

**Supplementary Figure S1.**

## STAR★METHODS

### KEY RESOURCES TABLE

The items in the key resources table (KRT) must also be reported alongside the description of their use in the method details section. Literature cited within the KRT must be included in the references list. Please **do not edit the headings or add custom headings or subheadings** to the KRT. We highly recommend using [RRIDs](https://scicrunch.org/resources) as the identifier for antibodies and model organisms in the KRT. To create the KRT, please use the template below or the [KRT webform](https://star-methods.com/). See the more detailed [Word table template](http://www.cell.com/pb-assets/journals/research/cell/methods/table-template1.docx) document for examples of how to list items.

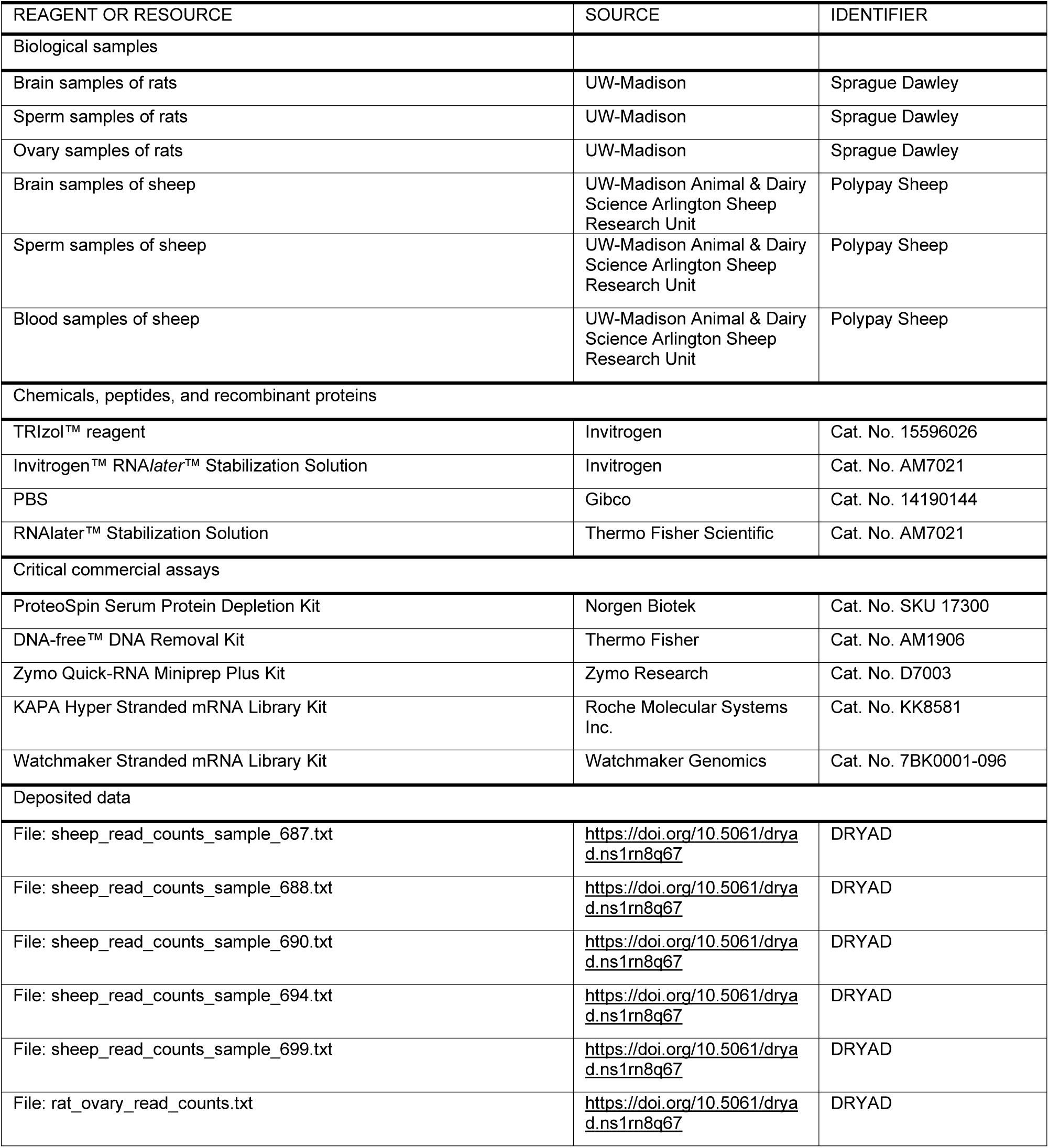
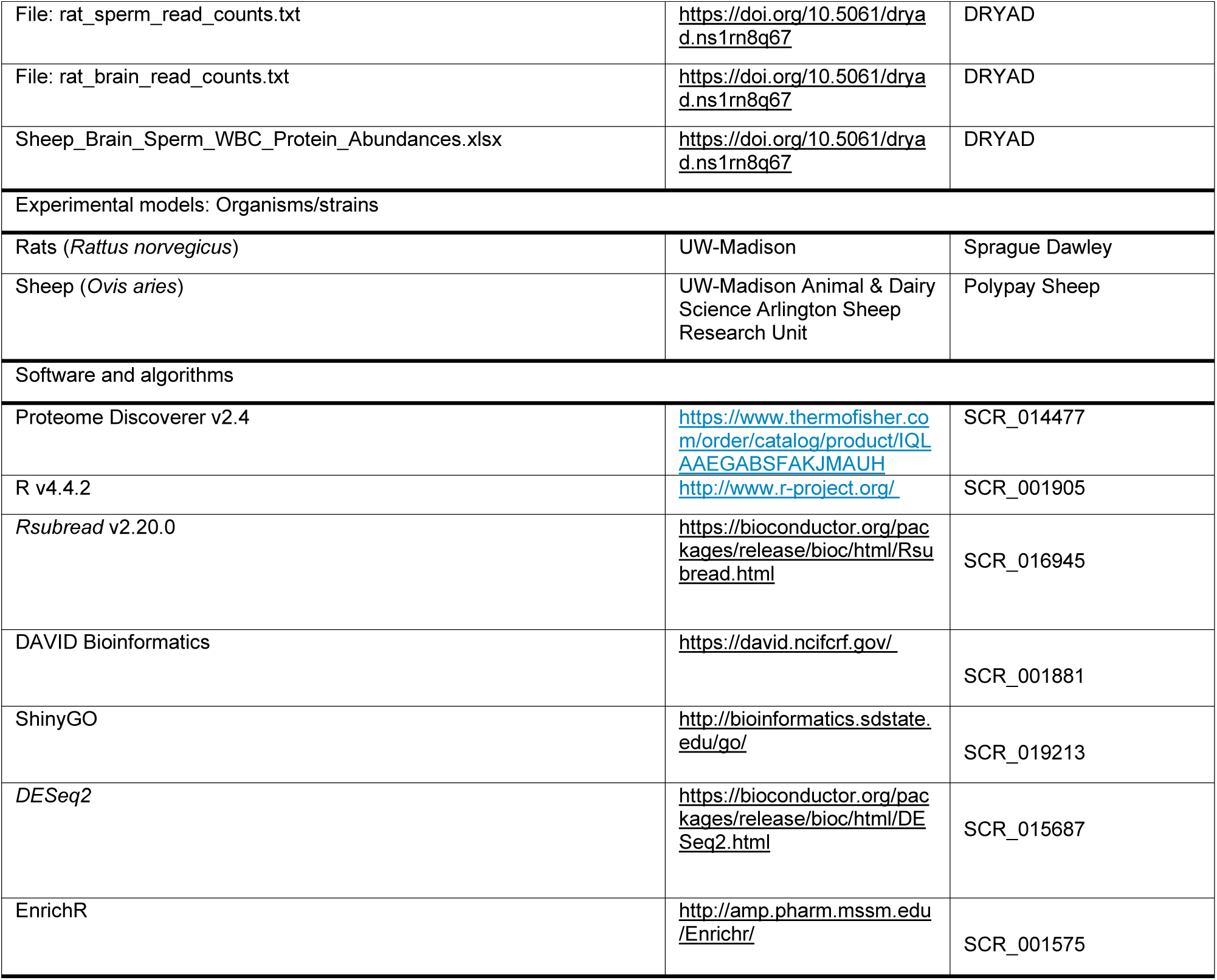

### EXPERIMENTAL MODEL AND STUDY PARTICIPANT DETAILS

A schematic representation of the experimental design detailing the sample collection, processing workflows, and analytical pipelines used in this study is shown in Figure 6.

**Figure 6.**
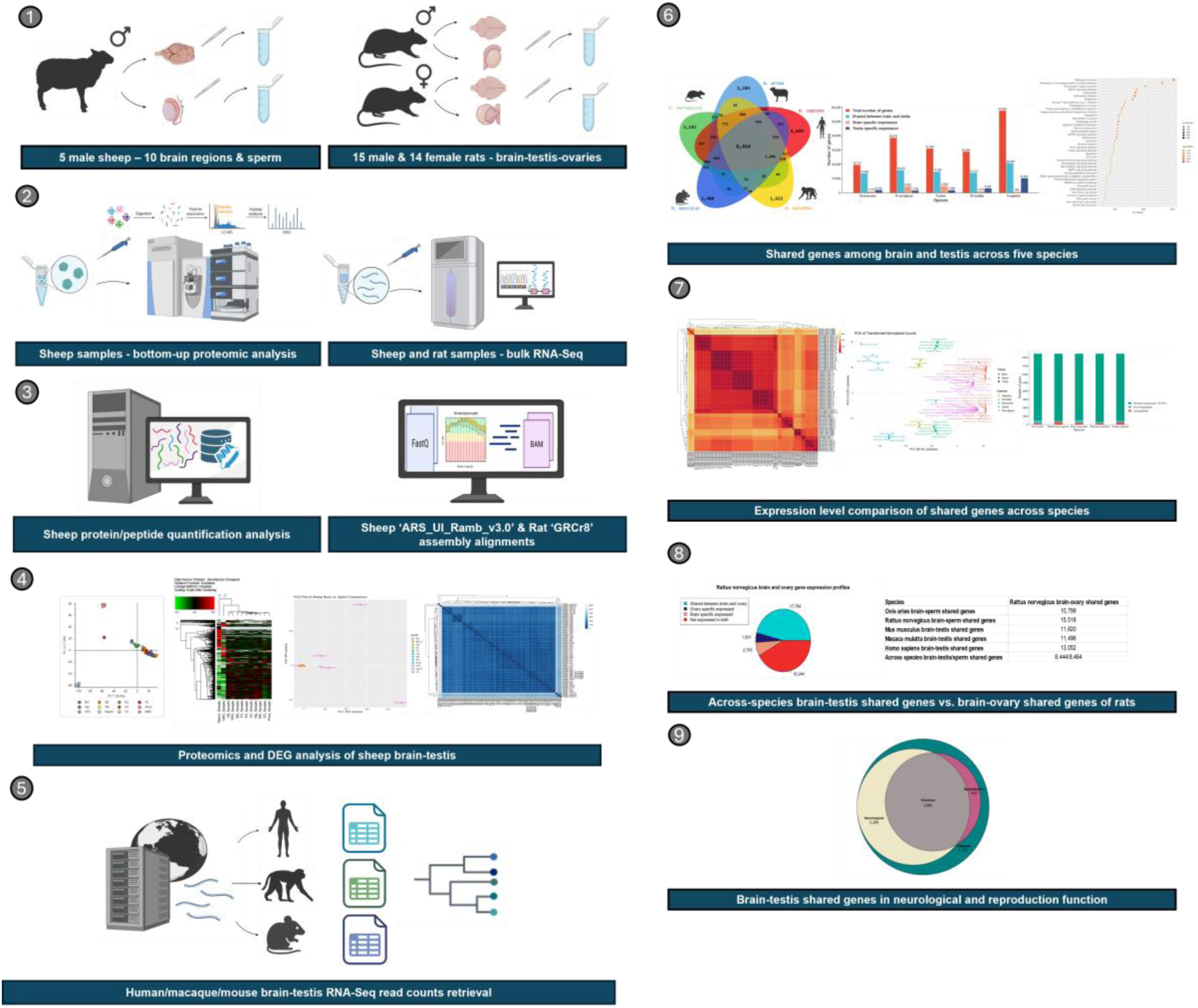
Schematic figure of experimental design.

All animal procedures were approved by the Institutional Animal Care and Use Committee (IACUC) of the University of Wisconsin–Madison (Protocol ID: A006488-R01) and conducted in accordance with institutional and national guidelines for the care and use of laboratory animals.

The experimental population comprised five post-pubertal Polypay rams (*Ovis aries*) raised under uniform environmental conditions, and 29 post-pubertal Sprague Dawley rats (*Rattus norvegicus*), including 15 males and 14 females.

## METHOD DETAILS

### Sample Collection in Rams

For proteomic and transcriptomic profiling, semen was collected from rams by electroejaculation using a Lane Pulsator V probe (Lane Manufacturing Inc., Denver, CO, USA). The ejaculate was collected in a 15-mL conical vial and immediately diluted with prewarmed (37 °C) semen extender (Minitube, Verona, WI).

Samples were centrifuged at 3000 × g for 10 min to pellet sperm, which were then washed twice with phosphate-buffered saline (PBS) and stored in RNAlater (Thermo Fisher Scientific, IL, USA) at −20 °C.

Blood samples from the same rams were collected in 5 mL EDTA-coated tubes to obtain white blood cells (WBC) for use as an outgroup in proteomic analyses, allowing comparison of sperm and WBC proteomes with those of various brain regions.

Rams were rendered unconscious by electrical stunning and exsanguinated at a commercial slaughter facility following standard industry practices. Heads were removed postmortem, placed on ice, and transported to a pathology facility for brain dissection. Whole brains were carefully extracted using a Stryker saw, sterile blades, forceps, and scissors. Each brain was divided into hemispheres, and the following regions were isolated: basal ganglia (BG), cerebellum (CB), entorhinal cortex (EC), frontal cortex (FC), hippocampus (HC), motor cortex (MC), prefrontal cortex (PC), pons, posterior parietal cortex (PPC), and visual cortex (VC). Tissue samples from each region were placed in either an empty 1.8 mL cryotube for flash-freezing or in a 1.8 mL cryotube containing ∼500 μL of RNAlater. All samples were flash-frozen in liquid nitrogen immediately after collection.

### Sample Collection in Rats

Whole brains and seminal fluid were collected from freshly euthanized male Sprague Dawley rats, while whole brains and ovaries were collected from females. Brains were extracted using sterile scissors and forceps, bisected into hemispheres, and placed either in RNAlater or in cryotubes without an additive. Ovaries were excised using sterile scissors and forceps. Seminal fluid was obtained by isolating the vas deferens and gently expressing the contents with the blunt end of sterile forceps into 1.8 mL cryotubes. All rat tissue and fluid samples were flash-frozen in liquid nitrogen immediately after collection.

### Proteomics

To assess the degree of molecular similarity among distinct brain regions and their overlap with sperm and WBC, we conducted comparative proteomic profiling across these tissues. Frozen sheep brain, sperm, and WBC samples were processed for proteome analysis beginning with the depletion of abundant proteins using the ProteoSpin Serum Protein Depletion Kit (Norgen Biotek, ON, Canada) following the manufacturer’s instructions. Protein concentrations were quantified at 280 nm using a NanoDrop One spectrophotometer (Thermo Fisher Scientific).

For in-solution digestion, 100 µg of total protein was used per sample. Each sample was supplemented with 1 µg of bovine serum albumin (BSA; Thermo Fisher Scientific) as an internal standard to ensure consistent quantification. Proteins were digested using trypsin/Lys-C (Promega, WI, USA) according to the manufacturer’s protocol. The resulting tryptic peptides were desalted and dried prior to LC-MS/MS analysis.

Dried peptides were reconstituted in 100 µL of mobile phase A (water with 0.1% formic acid), and 3 µg (3 µL) of peptides per sample were injected into the LC-MS/MS system. LC-MS/MS analyses were performed on a Dionex 3000 UHPLC system coupled to a Q-Exactive HF-X mass spectrometer (Thermo Fisher Scientific). Raw spectral data were searched against the *Ovis aries* UniProt reference proteome (ID: UP000009940) using Proteome Discoverer v2.4 (Thermo Fisher Scientific) as previously described ^36^.

Proteins with at least one detected peptide count were considered expressed, consistent with established mass spectrometry standards. Principal component analysis (PCA; Fig. 1A) and hierarchical clustering heatmap (Fig. 1B) were generated using protein abundances normalized to the spiked-in enolase standard. The heatmap was constructed based on Euclidean distance metrics.

### Transcriptomic Profiling

We next extended our analysis to transcriptomic profiling of distinct brain regions, focusing on the overlap between brain and sperm/testis gene expression across multiple species, including *Rattus norvegicus*, and on the shared transcriptomic profiles of brain and ovary tissues in *Rattus norvegicus*. For this purpose, we used same-hemisphere brain dissections and sperm samples from five rams, whole-brain and sperm samples from 15 male rats, and whole-brain and ovary samples from 14 female rats for total RNA extraction.

### RNA Extraction in Sheep

Approximately 25 mg of brain tissue preserved in RNAlater (Thermo Fisher Scientific) was homogenized in 1 mL of TRIzol reagent (Invitrogen, Waltham, MA, USA). Total RNA was extracted following the manufacturer’s protocol and treated with DNase I (Thermo Fisher Scientific) to remove residual genomic DNA prior to sequencing.

For sperm RNA extraction, approximately 200 µL of sperm preserved in RNAlater was diluted with 200 µL of PBS and centrifuged at 4000 rpm for 4 min to pellet sperm cells. The pellet was washed once in PBS, and the supernatant was discarded. To eliminate potential somatic cell contamination, the pellet was resuspended in 1 mL of somatic cell lysis buffer, incubated on ice for 4 min, and centrifuged again at 4000 rpm for 4 min. The resulting sperm pellet was resuspended in 1 mL of TRIzol reagent for total RNA extraction, followed by DNase I treatment to remove residual genomic DNA.

### RNA Extraction in Rats

RNA extraction from rat brain, ovary, and sperm samples followed a similar approach. Frozen brain samples were ground and homogenized in 1 mL of TRIzol reagent, followed by RNA extraction and DNase I treatment. Ovary and sperm samples were processed using the Zymo Quick-RNA Miniprep Plus Kit (Zymo Research, Irvine, CA, USA) to ensure high RNA yield and purity.

### RNA Sequencing

RNA sequencing was performed for all samples at the Roy J. Carver Biotechnology Center, University of Illinois at Urbana–Champaign, using the NovaSeq X Plus platform (Illumina, San Diego, CA, USA) with two 25B lanes (one per species) and V1.0 sequencing kits. Libraries were sequenced as 150 bp paired-end reads. *Ovis aries* RNA-Seq libraries were prepared with the KAPA Hyper Stranded mRNA Library Kit (Roche Molecular Systems Inc., Branchburg, NJ, USA), and *Rattus norvegicus* libraries with the Watchmaker Stranded mRNA Library Kit (Watchmaker Genomics, CO, USA). Libraries were pooled, quantified by qPCR, and sequenced for 151 cycles from both ends of each fragment.

Fastq files were generated and demultiplexed using bcl2fastq v2.20 for *Ovis aries* and v4.1.17 for *Rattus norvegicus* (Illumina Inc.). On average, 122,228,640 reads were generated per sheep sample and 122,551,952 reads per rat sample. Read quality was assessed with MultiQC, and adapter trimming and quality filtering (ASCII offset 33) were performed using Trimmomatic v0.39 ^37,38^. Alignment was carried out with STAR v2.7.5a ^39^ against the *Rattus norvegicus* reference genome GRCr8 (GCF_036323735.1) and the *Ovis aries* reference genome ARS-UI_Ramb_v3.0 (GCA_016772045.2), using their corresponding annotation files. Alignment rates exceeded 80% for most samples. Read counts were obtained in R (v4.4.2) using the *Rsubread* package (v2.20.0) ^40,41^. Principal component analysis (PCA; Fig. 1C) and hierarchical clustering heatmaps (Fig. 1D) of normalized read counts were generated to assess transcriptomic similarities and distances among tissues.

### RNA-Seq Data Retrieval

To expand the evolutionary scope of our analysis across species, brain and testis RNA-Seq read count data from *Mus musculus*, *Macaca mulatta*, and *Homo sapiens* were retrieved from publicly available databases.

Brain and testis transcriptomic profiles from 15 adult male *Homo sapiens* (aged 50–59 years) who died of natural causes with no known disease history were obtained from the Genotype-Tissue Expression (GTEx) database as non-normalized read counts ^42^.

To ensure inclusion of only healthy animals without reproductive or neurological disorders, brain and testis RNA-Seq read count data from nine post-pubertal male *Mus musculus* samples were obtained from the NCBI Gene Expression Omnibus (GEO) under accession numbers GSE194203, GSE271865, and GSE194203 ^43^.

For *Macaca mulatta*, only three matched brain and testis RNA-Seq datasets from healthy rhesus macaques were available; these were retrieved from the EMBL-EBI Expression Atlas under accession number E-MTAB-2799 ^44^.

Gene annotations varied across species, leading to different numbers of protein-coding and non-coding genes represented in each dataset. Notably, the *Mus musculus* datasets contained 19,773 annotated protein-coding genes, which may have constrained the number of genes identified in subsequent comparative analyses.

### Identification of Brain–Sperm/Testis Shared Genes

To identify genes with shared expression between the brain and sperm/testis within and across species, we conducted a comparative shared-expression analysis followed by cross-species gene intersection. The analysis included RNA-Seq read count data from five species: the experimentally generated brain and sperm datasets for *Ovis aries* and *Rattus norvegicus*, and publicly available brain and testis RNA-Seq datasets for *Mus musculus*, *Macaca mulatta*, and *Homo sapiens*.

A uniform threshold was applied to define gene expression across all species: genes were considered expressed if they exhibited ≥10 read counts in at least half of the samples within a tissue. For datasets employing gene identifiers other than the official gene symbol, ID conversion was performed using the DAVID Bioinformatics Resources ID Conversion Tool ^45^. After extracting the sets of genes expressed in both brain and sperm/testis for each species, we overlapped these gene lists to identify the conserved set of brain–sperm/testis-shared genes across species.

Functional enrichment analysis of the conserved gene set was then performed using ShinyGO v0.82 ^46^, applying a false discovery rate (FDR) correction (q < 0.05) and retaining pathways containing 2-1000 genes. The top 40 Kyoto Encyclopedia of Genes and Genomes (KEGG) pathways and Gene Ontology (GO) biological processes were reported as significantly enriched.

A summary of these findings is presented in Figure 2, including: (A) the Venn diagram of shared genes between brain and sperm/testis across all species, (B) the cross-species distribution of annotated genes, (C) enriched KEGG pathways, and (D) enriched GO biological processes.

### Tissue Enrichment Expression Analysis of Shared Genes

To evaluate the conservation status of shared genes in terms of “similar expression” versus “tissue-enriched” expression in brain and sperm/testis across species, we performed a tissue enrichment expression analysis. Although no universal threshold exists for defining tissue enrichment, previous studies have applied fold-change cutoffs ranging from 2– to 50-fold depending on the research question and desired stringency ^47–49^.

Given that our dataset had already been refined to 8,464 genes shared across species, and subtle expression differences were not the focus of this analysis, we applied a stringent criterion: genes exhibiting ≥10-fold differential expression with a false discovery rate (FDR)-corrected q-value < 0.05 were classified as “tissue-enriched,” while those below this threshold were considered “similarly expressed.” Differential expression analysis between brain and sperm/testis within each species was performed using the *DESeq2* R package, with read counts normalized using the median-of-ratios method ^50^. The resulting distributions of gene expression patterns are illustrated in Fig. 3.

### Overlap Between Cross-Species Brain–Sperm/Testis and Brain–Ovary Shared Genes in Rattus norvegicus

To examine the relationship between cross-species brain–sperm/testis-shared genes and brain–ovary-shared genes in *Rattus norvegicus*, we performed a shared expression analysis using experimentally obtained brain and ovary transcriptomes from 14 female rats. Genes with ≥10 read counts in at least seven samples per tissue were considered expressed. The expressed gene lists from the brain and the ovary were then intersected to identify brain-specific, ovary-specific, and shared genes. The results of this analysis, including the overlap with cross-species brain–sperm/testis shared genes, are presented in Fig. 4.

### Identification of Genes Involved in Nervous and Reproductive System Functions

To identify genes associated with nervous system and reproductive functions, we performed a comprehensive functional annotation analysis using ten databases: Jensen’s Diseases Curated dataset, Human Phenotype Ontology (HPO), Mouse Genome Informatics (MGI), Kyoto Encyclopedia of Genes and Genomes (KEGG), WikiPathways, Gene Association Database (GAD), Online Mendelian Inheritance in Man (OMIM), DisGeNET, UniProt Keywords, and Gene Ontology (Biological Processes). Annotations were accessed through bioinformatic tools including DAVID Bioinformatics, EnrichR, and ShinyGO ^46,51–53^.

Following annotation of brain–sperm/testis shared genes across these databases, Microsoft’s CoPilot AI was used to classify genes into four categories: (1) annotated only with neurological functions, (2) annotated only with reproductive functions, (3) annotated with both neurological and reproductive functions, and (4) uncharacterized in either context. This classification was performed by providing the AI with annotation lists in manageable chunks and instructing it to systematically query each database, match gene annotations, interpret annotation descriptions, and assign each gene to the appropriate functional category ^54^.

## QUANTIFICATION AND STATISTICAL ANALYSIS

All quantification and statistical analyses in this study were mentioned in their relevant sections at “Method details”.

Briefly, proteins with a minimum of one detected peptide count were considered ‘expressed’, consistent with established mass spectrometry standards. Principal component analysis (PCA; Fig. 1A) and hierarchical clustering heatmap (Fig. 1B) were generated using protein abundances normalized to the spiked-in enolase standard. The heatmap was constructed based on Euclidean distance metrics. Both were generated using Proteome Discoverer v2.4 (Thermo Fisher Scientific). All the remaining quantification and statistical analysis were performed using R environment (v4.4.2) with various packages as detailed in “Method details” ^41^. Read counts were obtained using the *Rsubread* package (v2.20.0) ^40^. Principal component analysis (PCA; Fig. 1C) and hierarchical clustering heatmaps (Fig. 1D) of normalized read counts were generated to assess transcriptomic similarities and distances among tissues. A uniform threshold was applied to define gene expression across all species: genes were considered expressed if they exhibited ≥10 read counts in at least half of the samples within a tissue. Functional enrichment analysis of the conserved gene set was then performed using ShinyGO v0.82 ^46^, applying a false discovery rate (FDR) correction (q < 0.05) and retaining pathways containing 2-1000 genes. Genes exhibiting ≥10-fold differential expression with a false discovery rate (FDR)-corrected q-value < 0.05 were classified as “tissue-enriched,” while those below this threshold were considered “similarly expressed.” Differential expression analysis between brain and sperm/testis within each species was performed using the *DESeq2* R package, with read counts normalized using the median-of-ratios method ^50^. The resulting distributions of gene expression patterns are illustrated in Fig. 3.

## REFERENCES

1. Strickberger, M.W. (2000). Evolution 3rd Edition. (Jones & Bartlett Learning).

2. Cole, E.F., Morand-Ferron, J., Hinks, A.E., and Quinn, J.L. (2012). Cognitive Ability Influences Reproductive Life History Variation in the Wild. Current Biology 22, 1808–1812. 10.1016/j.cub.2012.07.051.

3. Keagy, J., Savard, J.F., and Borgia, G. (2009). Male satin bowerbird problem-solving ability predicts mating success. Anim Behav 78, 809–817. 10.1016/j.anbehav.2009.07.011.

4. Morand-Ferron, J., and Quinn, J.L. (2015). The evolution of cognition in natural populations. Trends Cogn Sci 19, 235–237. 10.1016/j.tics.2015.03.005.

5. Arden, R., Gottfredson, L.S., Miller, G., and Pierce, A. (2009). Intelligence and semen quality are positively correlated. Intelligence 37, 277–282. 10.1016/j.intell.2008.11.001.

6. Duboule, D., and Wilkins, A.S. (1998). The evolution of ‘bricolage’. Trends in Genetics 14, 54–59. 10.1016/S0168-9525(97)01358-9.

7. Wilda, M., Bächner, D., Zechner, U., Kehrer-Sawatzki, H., Vogel, W., Hameister, H., and Humangenetik, A. (2000). Do the constraints of human speciation cause expression of the same set of genes in brain, testis, and placenta? Cytogenetics and cell genetics 91, 300–302. 10.1159/000056861

8. Yan, J., Zhang, H., Liu, Y., Zhao, F., Zhu, S., Xie, C., Tang, T.S., and Guo, C. (2016). Germline deletion of huntingtin causes male infertility and arrested spermiogenesis in mice. J Cell Sci 129, 492–501. 10.1242/jcs.173666.

9. Crawford, D.C., Acuña, J.M., and Sherman, S.L. (2001). FMR1 and the fragile X syndrome: Human genome epidemiology review. Genetics in Medicine 3, 359–371. 10.1097/00125817-200109000-00006.

10. Man, L., Lekovich, J., Rosenwaks, Z., and Gerhardt, J. (2017). Fragile X-Associated Diminished Ovarian Reserve and Primary Ovarian Insufficiency from Molecular Mechanisms to Clinical Manifestations. Front Mol Neurosci Volume 10,290. 10.3389/fnmol.2017.00290.

11. Slegtenhorst-Eegdeman, K.E., de Rooij, D.G., Verhoef-Post, M., van de Kant, H.J.G., Bakker, C.E., Oostra, B.A., Grootegoed, J.A., and Themmen, A.P.N. (1998). Macroorchidism in FMR1 Knockout Mice Is Caused by Increased Sertoli Cell Proliferation during Testicular Development*. Endocrinology 139, 156–162. 10.1210/endo.139.1.5706.

12. Guo, J., Zhu, P., Wu, C., Yu, L., Zhao, S., and Gu, X. (2003). In silico analysis indicates a similar gene expression pattern between human brain and testis. Cytogenet Genome Res 103, 58–62. 10.1159/000076290.

13. Matos, B., Publicover, S.J., Castro, L.F.C., Esteves, P.J., and Fardilha, M. (2021). Brain and testis: More alike than previously thought? Preprint at Royal Society Publishing, 10.1098/rsob.200322.

14. Erdö, S.L., and Wekerle, L. (1990). GABAA type binding sites on membranes of spermatozoa. Life Sci 47, 1147–1151. 10.1016/0024-3205(90)90175-Q.

15. Ritta, M.N., Calamera, J.C., and Bas, D.E. (1998). Occurrence of GABA and GABA receptors in human spermatozoa. Mol Hum Reprod 4, 769–773. 10.1093/molehr/4.8.769.

16. Meizel, S. (2004). The sperm, a neuron with a tail: “Neuronal” receptors in mammalian sperm. Preprint, 10.1017/S1464793103006407.

17. Kaewman, P., Nudmamud-Thanoi, S., Amatyakul, P., and Thanoi, S. (2021). High mRNA expression of GABA receptors in human sperm with oligoasthenoteratozoospermia and teratozoospermia and its association with sperm parameters and intracytoplasmic sperm injection outcomes. Clin Exp Reprod Med 48, 50–60. 10.5653/cerm.2020.03972.

18. Gross, N., Taylor, T., Crenshaw, T., and Khatib, H. (2020). The Intergenerational Impacts of Paternal Diet on DNA Methylation and Offspring Phenotypes in Sheep. Front Genet Volume 11-2020.10.3389/fgene.2020.597943.

19. Townsend, J., Braz, C.U., Taylor, T., and Khatib, H. (2023). Effects of paternal methionine supplementation on sperm DNA methylation and embryo transcriptome in sheep. Environ Epigenet 9, dvac029. 10.1093/eep/dvac029.

20. Braz, C.U., Passamonti, M.M., and Khatib, H. (2024). Characterization of genomic regions escaping epigenetic reprogramming in sheep. Environ Epigenet 10. 10.1093/eep/dvad010.

21. Kizilaslan, M., Braz, C.U., Townsend, J., Taylor, T., Crenshaw, T.D., and Khatib, H. (2025). Transgenerational Epigenetic and Phenotypic Inheritance Across Five Generations in Sheep. Int J Mol Sci 26. 10.3390/ijms26136412.

22. Doronina, L., Reising, O., Clawson, H., Churakov, G., and Schmitz, J. (2022). Euarchontoglires Challenged by Incomplete Lineage Sorting. Genes (Basel) 13. 10.3390/genes13050774.

23. Senger, P.L. (2012). Pathways to pregnancy and parturition 3rd edition. Redmond OR: Current Conceptions.

24. Mila, M., Alvarez-Mora, M.I., Madrigal, I., and Rodriguez-Revenga, L. (2018). Fragile X syndrome: An overview and update of the FMR1 gene. Preprint at Blackwell Publishing Ltd, 10.1111/cge.13075.

25. Dijkstra, A.A., Haify, S.N., Verwey, N.A., Prins, N.D., van der Toorn, E.C., Rozemuller, A.J.M., Bugiani, M., den Dunnen, W.F.A., Todd, P.K., Charlet-Berguerand, N., et al. (2021). Neuropathology of FMR1-premutation carriers presenting with dementia and neuropsychiatric symptoms. Brain Commun 3. 10.1093/braincomms/fcab007.

26. Pastore, L.M., and Johnson, J. (2014). The FMR1 gene, infertility, and reproductive decision-making: A review. Preprint at Frontiers Research Foundation, 10.3389/fgene.2014.00195.

27. Stelzer, G., Rosen, N., Plaschkes, I., Zimmerman, S., Twik, M., Fishilevich, S., Stein, T.I., Nudel, R., Lieder, I., Mazor, Y., et al. (2016). The GeneCards Suite: From Gene Data Mining to Disease Genome Sequence Analyses. Curr Protoc Bioinformatics 54, 1.30.1–1.30.33. 10.1002/cpbi.5.

28. Stephenson, J.R., Wang, X., Perfitt, T.L., Parrish, W.P., Shonesy, B.C., Marks, C.R., Mortlock, D.P., Nakagawa, T., Sutcliffe, J.S., and Colbran, R.J. (2017). A novel human CAMK2a mutation disrupts dendritic morphology and synaptic transmission, and causes ASD-related behaviors. Journal of Neuroscience 37, 2216–2233. 10.1523/JNEUROSCI.2068-16.2017.

29. Shabtay, O., and Breitbart, H. (2016). CaMKII prevents spontaneous acrosomal exocytosis in sperm through induction of actin polymerization. Dev Biol 415, 64–74. 10.1016/j.ydbio.2016.05.008.

30. Tatone, C., Delle Monache, S., Iorio, R., Caserta, D., Di Cola, M., and Colonna, R. (2002). Possible role for Ca 2 calmodulin-dependent protein kinase II as an effector of the fertilization Ca 2 signal in mouse oocyte activation. Molecular human reproduction, 8, 750–757. 10.1093/molehr/8.8.750

31. Kaneko, U., Toyabe, S.I., Hara, M., and Uchiyama, M. (2006). Increased mutations of CD72 transcript in B-lymphocytes from adolescent patients with systemic lupus erythematosus. Pediatric Allergy and Immunology 17, 565–571. 10.1111/j.1399-3038.2006.00466.x.

32. Tian, L., Wang, Y., Zhang, Z., Feng, X., Xiao, F., and Zong, M. (2023). CD72, a new immune checkpoint molecule, is a novel prognostic biomarker for kidney renal clear cell carcinoma. Eur J Med Res 28. 10.1186/s40001-023-01487-8.

33. Dodeller, F., Gottar, M., Huesken, D., Iourgenko, V., and Cenni, B. (2008). The Lysosomal Transmembrane Protein 9B Regulates the Activity of Inflammatory Signaling Pathways*. Journal of Biological Chemistry 283, 21487–21494. 10.1074/jbc.M801908200.

34. Tracey, K.J. (2015). Approaching the next revolution? Evolutionary integration of neural and immune pathogen sensing and response. Cold Spring Harb Perspect Biol 7. 10.1101/cshperspect.a016360.

35. Baluška, F., Miller, W.B., and Reber, A.S. (2023). Cellular and evolutionary perspectives on organismal cognition: From unicellular to multicellular organisms. Preprint at Oxford University Press, 10.1093/biolinnean/blac005.

36. Townsend, J., Peng, Z., Kizilaslan, M., Taylor, T., Yang, Z., Ahsan, N., and Khatib, H. (2025). Methionine supplementation-induced alteration of sheep seminal plasma miRNAs and proteome. J Anim Sci 103. 10.1093/jas/skaf192.

37. Bolger, A.M., Lohse, M., and Usadel, B. (2014). Trimmomatic: A flexible trimmer for Illumina sequence data. Bioinformatics 30, 2114–2120. 10.1093/bioinformatics/btu170.

38. Ewels, P., Magnusson, M., Lundin, S., and Käller, M. (2016). MultiQC: Summarize analysis results for multiple tools and samples in a single report. Bioinformatics 32, 3047–3048. 10.1093/bioinformatics/btw354.

39. Dobin, A., Davis, C.A., Schlesinger, F., Drenkow, J., Zaleski, C., Jha, S., Batut, P., Chaisson, M., and Gingeras, T.R. (2013). STAR: Ultrafast universal RNA-seq aligner. Bioinformatics 29, 15–21. 10.1093/bioinformatics/bts635.

40. Liao, Y., Smyth, G.K., and Shi, W. (2019). The R package Rsubread is easier, faster, cheaper and better for alignment and quantification of RNA sequencing reads. Nucleic Acids Res 47. 10.1093/nar/gkz114.

41. R Core Team (2024). R: A Language and Environment for Statistical Computing, Vienna, Austria. https://www.R-project.org/.

42. Lonsdale, J., Thomas, J., Salvatore, M., Phillips, R., Lo, E., Shad, S., Hasz, R., Walters, G., Garcia, F., Young, N., et al. (2013). The Genotype-Tissue Expression (GTEx) project. Preprint, 10.1038/ng.2653.

43. Barrett, T., Wilhite, S.E., Ledoux, P., Evangelista, C., Kim, I.F., Tomashevsky, M., Marshall, K.A., Phillippy, K.H., Sherman, P.M., Holko, M., et al. (2013). NCBI GEO: Archive for functional genomics data sets-Update. Nucleic Acids Res 41. 10.1093/nar/gks1193.

44. Merkin, J., Russell, C., Chen, P., and Burge, C.B. (2012). Evolutionary Dynamics of Gene and Isoform Regulation in Mammalian Tissues. Science (1979) 338, 1593–1599. 10.1126/science.1228186.

45. Sherman, B.T., Stephens, R., Baseler, M.W., Lane, H.C., and Lempicki, R.A. (2008). DAVID gene ID conversion tool. Bioinformation 2, 428.

46. Ge, S.X., Jung, D., and Yao, R. (2020). ShinyGO: a graphical gene-set enrichment tool for animals and plants. Bioinformatics 36, 2628–2629. 10.1093/bioinformatics/btz931.

47. Fagerberg, L., Hallström, B.M., Oksvold, P., Kampf, C., Djureinovic, D., Odeberg, J., Habuka, M., Tahmasebpoor, S., Danielsson, A., Edlund, K., et al. (2014). Analysis of the Human Tissue-specific Expression by Genome-wide Integration of Transcriptomics and Antibody-based Proteomics*. Molecular & Cellular Proteomics 13, 397–406. 10.1074/mcp.M113.035600.

48. Rawal, H.C., Angadi, U., and Mondal, T.K. (2021). TEnGExA: An R package-based tool for tissue enrichment and gene expression analysis. Brief Bioinform 22. 10.1093/bib/bbaa221.

49. Ryaboshapkina, M., and Hammar, M. (2019). Tissue-specific genes as an underutilized resource in drug discovery. Sci Rep 9. 10.1038/s41598-019-43829-9.

50. Love, M.I., Huber, W., and Anders, S. (2014). Moderated estimation of fold change and dispersion for RNA-seq data with DESeq2. Genome Biol 15, 550. 10.1186/s13059-014-0550-8.

51. Huang, D.W., Sherman, B.T., and Lempicki, R.A. (2009). Systematic and integrative analysis of large gene lists using DAVID bioinformatics resources. Nat Protoc 4, 44–57. 10.1038/nprot.2008.211.

52. Chen, E.Y., Tan, C.M., Kou, Y., Duan, Q., Wang, Z., Meirelles, G.V., Clark, N.R., and Ma’ayan, A. (2013). Enrichr: interactive and collaborative HTML5 gene list enrichment analysis tool. BMC Bioinformatics 14, 128. 10.1186/1471-2105-14-128.

53. Sherman, B.T., Hao, M., Qiu, J., Jiao, X., Baseler, M.W., Lane, H.C., Imamichi, T., and Chang, W. (2022). DAVID: a web server for functional enrichment analysis and functional annotation of gene lists (2021 update). Nucleic Acids Res 50, W216–W221. 10.1093/nar/gkac194.

54. Microsoft (2025). Copilot: AI-powered orchestration layer built on foundational models. https://copilot.microsoft.com/

