## Supplementary Figure 1 for "Across Species Identification of Genes Bridging Cognition and Reproduction"

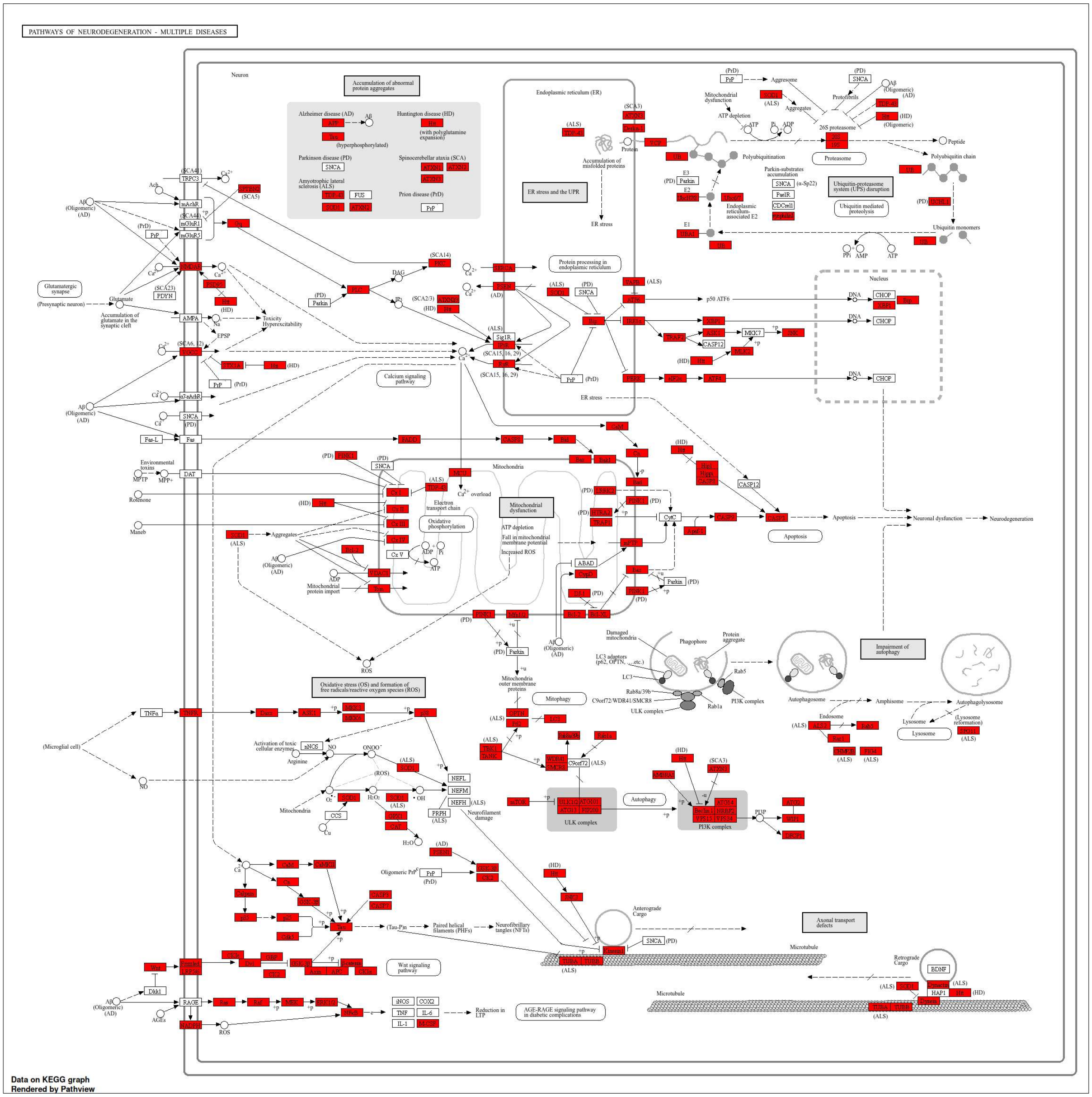


**Supplementary Fig. S1.** Pathways of neurodegeneration presented by Kyoto Encyclopedia of Genes and Genomes. Red-marked genes are the genes found to be shared between the brain and sperm/testis across species.
